# A compact breast shape acquisition system for improving diffuse optical tomography image reconstructions

**DOI:** 10.1101/2022.11.20.517255

**Authors:** Morris Vanegas, Miguel Mireles, Edward Xu, Shijie Yan, Qianqian Fang

**Affiliations:** Department of Bioengineering, Northeastern University, 02115 Boston, USA; Department of Electrical and Computer Engineering, Northeastern University, 02115, USA

## Abstract

Diffuse optical tomography (DOT) has been investigated for diagnosing malignant breast lesions but its accuracy relies on model-based image reconstructions which in turn depends on the accuracy of breast shape acquisition. In this work, we have developed a dual-camera structured light imaging (SLI) breast shape acquisition system tailored for a mammography-like compression setting. Illumination pattern intensity is dynamically adjusted to account for skin tone differences while thickness-informed pattern masking reduces artifacts due to specular reflections. This compact system is affixed to a rigid mount that can be installed into existing mammography or parallel-plate DOT systems without the need for camera-projector re-calibration. Our SLI system produces sub-millimeter resolution with a mean surface error of 0.26 mm. This breast shape acquisition system results in more accurate surface recovery, with an average 1.6-fold reduction in surface estimation errors over a reference method via contour extrusion. Such improvement translates to 25% to 50% reduction in mean squared error in the recovered absorption coefficient for a series of simulated tumors 1-2 cm below the skin.

## 1. Introduction

Breast cancer is the most commonly diagnosed cancer in women worldwide with an estimated 1,918,030 new cases in 2022 in the United States alone [1]. X-ray mammography is the primary breast cancer screening technique [2] used for early detection to reduce mortality rates [3]. Despite its recommendation for screening, x-ray mammography suffers from a high false-positive diagnostic rate [3, 4]. The technique lacks both strong structural contrast between healthy and tumor tissue and the ability to quantify tissue functions to assess benign versus malignancy [5]. These limitations have led researchers to investigate using diffuse optical tomography (DOT) techniques to characterize the breast tumor’s physiology to lower false-positive diagnoses.

Unlike x-ray mammography, DOT is an imaging modality that uses non-ionizing near-infrared (NIR) radiation to yield three-dimensional (3-D) maps of the optical properties of illuminated tissue [6–9]. Biological tissues’ primary absorbers in the spectral window from around 600 to 1000 nm have relatively low absorption, allowing NIR light to penetrate through centimeters of tissues [10]. This allows the quantification of physiological properties such as hemoglobin concentration, blood volume, and blood oxygen saturation [5, 6]. Malignant tumors generally demand a greater blood supply than their surrounding tissues, leading to increased light absorption that can be delineated using spectroscopy and imaging methods, making DOT particularly useful for breast cancer imaging diagnosis [11–15]. Additionally, the low spatial resolution of DOT [16] can be improved by a multi-modal approach with x-ray mammography [17–20]. DOT images are known for low spatial resolution largely caused by the high scattering properties of biological tissues [6]. The high scattering present in the breast tissue redirects photons to traverse large overlapping probing volumes before their detection, greatly reducing the locality of the measurements and resulting in blurry images. Mathematically, this is reflected as the severe ill-posedness of the inverse problem. Parallel-plate compression of breast tissues has been used in an x-ray mammography scan to minimize overlapping tissues and has also been explored for a number of standalone [14, 21] and multi-modal DOT breast imaging systems [20, 22, 23]. Obtaining breast surface information to aid quantitative analysis of imaging data has garnered interest from a number of applications, including digital breast tomosynthesis (DBT) [24] and magnetic resonance imaging (MRI) scans [25, 26].

For multi-modal DOT systems, the 3-D shape of the breast can be estimated using the structural imaging modality such as DBT [27] or MRI [28]. When a 3-D imaging modality is not available, two-dimensional (2-D) mammography [19] has also been used to provide the shape information. In such case, a simple way to recover a 3-D breast surface is to extrude the 2-D breast contour along the compression axis [29, 30], or sweep the 2-D breast contour along the contour line extracted from an orthogonal view [31]. These methods either rely on assumptions about the 3-D location of certain features (e.g. mamilla position) or assume a constant curvature of the breast along the sweeping direction. For more accurate reconstructions of tissue optical properties, especially near the surface, measuring 3-D breast surface accurately can be greatly beneficial.

Accurately acquiring breast 3-D surface shapes has gained clinical acceptance due in large part to the plastic and reconstructive surgery communities [32, 33]. The two prominent techniques for 3-D breast surface imaging are stereophotogrammetry and laser scanning [34]. Stereophotogrammetry works by overlaying multiple images of an object taken from different view angles and triangulating the location of the object based on matching features in the multiple images [35–37]. In addition to requiring multiple cameras to increase accuracy [38], this technique is also heavily influenced by lighting conditions since features need to be extracted from multiple viewpoints [39]. Another limitation is the “ptosis error” seen during scanning of ptotic or larger breasts [40]. This arises due to the small field of view of stereophotogrammetry systems, leading to inaccuracies in breast surface and volume estimations due to the portions of the breast that are not illuminated. Laser scanning is a surface imaging technique in which rays from a laser beam are reflected off an object and detected by a detector [41]. Although laser-based acquisition systems can produce more accurate surfaces [42], these systems tend to be expensive [43, 44] and require the need for very precise setups [45]. Recently, the use of patterned-lasers and orientation-sensitive detectors has led to an increase in portable 3-D laser scanning devices [46]. While low-cost laser-based depth sensors have been widely deployed in game consoles such as Xbox or PlayStation, they are only designed to achieve relatively low spatial accuracy compared to dedicated 3-D scanners. Still, patterned-laser-based surface acquisition systems generally require a minimum scanner-to-target distance of 35 cm [47, 48]. Additionally, their typical housing is too large to fit between mammography compression plates [25, 48]. Bulky instrumentation and long minimum working distance requirements make stereophotogrammetry and laser scanning techniques infeasible in the confined, low-light mammography setting.

Another widely used 3-D surface acquisition technique is structured light imaging (SLI) [49,50]. SLI works by illuminating an object with two-dimensional spatially varying patterns with a projector and capturing images from the illuminated object using cameras [51]. The distortion of the specially designed patterns provides information regarding the shape of the object. Calibration of the camera-projector system is easily done by capturing images of a known planar pattern (e.g. a checkerboard). With the ability to use off-the-shelf components, a simple setup with a single projector and camera, and a robust and simple calibration method, SLI is positioned to provide accurate, fast, and cost-effective breast surface scanning [49]. However, similar to most patterned-laser surface scanners, commercially available SLI systems have long minimal working distance requirements and large assemblies that cannot readily fit within the confined mammography compression plates [50, 52].

In this work, we have developed a low-profile dual-camera SLI breast shape acquisition system specifically tailed for use in the confined space between parallel breast compression plates. This system can be incorporated with standalone DOT breast scanners or multi-modal DOT systems combined with mammography or DBT, with a minimal scanner-to-target distance between 10 and 15 cm. In the following sections, we first describe the design of the SLI breast scanner and detail the methods for adaptive illumination for subject-specific skin tones as well as approaches to reduce specular reflection from the compression plates. We then compare several traditional surface acquisition methods that leverage mammography images against our SLI-based breast surface acquisition system and quantify the impact of improved breast surface acquisition on the recovery of breast lesions using a series of simulations.

## 2. Methods

Here, we first briefly describe a mammography-mimicking parallel-plate transmission breast DOT system, designed to be used as either a standalone DOT scan of a compressed breast or combined with separately acquired mammography or DBT images in a multi-modality data analysis [18, 53]. Then we elaborate on the SLI breast surface acquisition sub-system and surface data processing pipeline. This custom SLI system is designed to be compact, low-cost, and specifically tailored towards the low-light and space-confined mammography compression settings. Next, we compare this approach with a number of alternative breast shape acquisition methods, including single-axis and dual-axis contour-line extrusion methods, and create benchmarks to quantify the surface accuracy improvement of the proposed method. Finally, we compare results from a series of DOT image reconstructions based on simulated data with various surface acquisition methods to further quantify the impact of breast surface accuracy, especially for the accurate recovery of tumors embedded at various depths.

### 2.1. Wide-field parallel-plate transmission breast DOT system

A parallel-plate transmission optical tomography system with the capability of imaging a breast with mammography-mimicking compression was built where the proposed SLI system is embedded between the compression plate to provide accurate measurement of the breast surface [Fig. 1(c)]. The breast is compressed by a pair of acrylic plates, with one plate mounted at the stationary end of a linear stage (BiSlide, Velmex, Bloomfield, NY, USA). An acrylic mammography compression paddle is mounted on the moving stage of the linear stage, allowing for a plate separation ranging from 300 mm (fully released) to 0 mm (fully closed) using a 2-phase stepper motor (Oriental Motors, Braintree, MA, USA). A linear encoder (ETI Systems, Carlsbad, CA, USA) is connected between the pair of compression plates to measure their separation. The entire breast compression assembly is mounted on a rotatory table (Lintech, Monrovia, CA, USA), controlled by a foot paddle to permit mammography-like lateral-oblique compression. This breast DOT design specifically enables registration of structural information from separately acquired mammography scans with the DOT images using the methods detailed in our previous studies [18]. The details of this breast DOT system will be described in a separate publication.

**Fig. 1.**
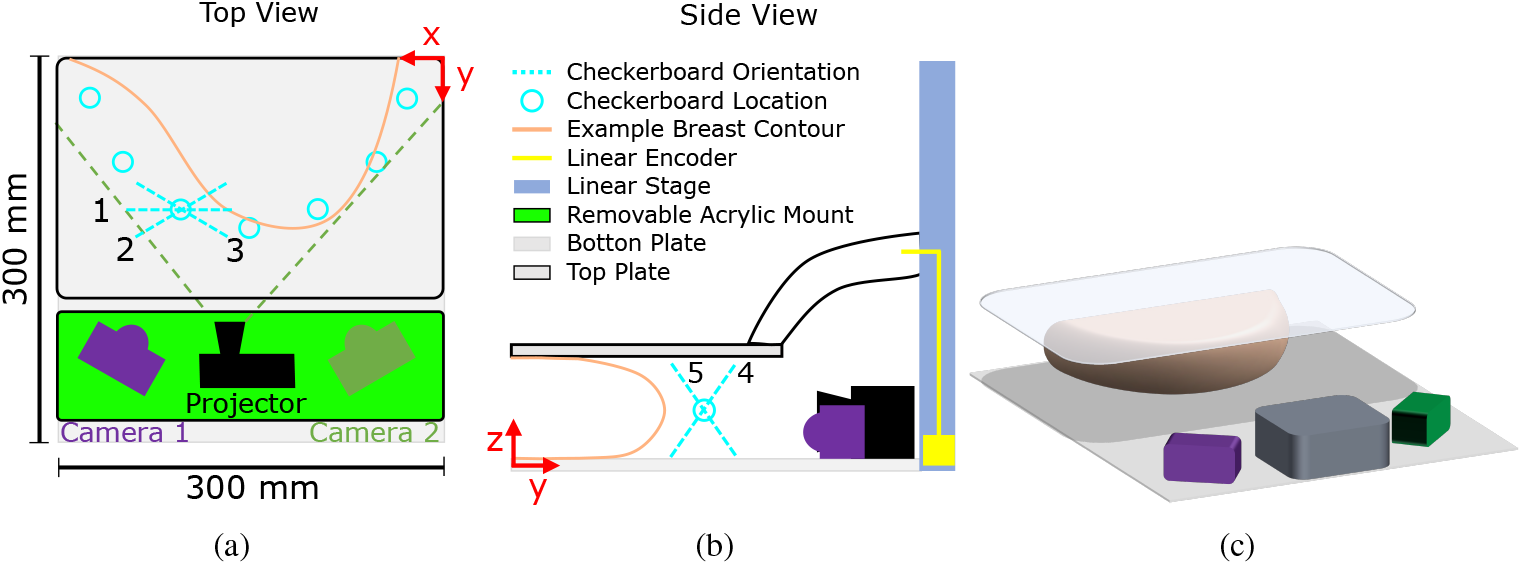
(a) Top-view of the breast compression compartment – upper: SLI system; bottom: horizontal cross-section (orange line) of the compressed breast with blue circles indicating the placement of the checkerboard used for system calibration. Numbers 1-5 indicate the 5 board orientations repeated at each location for calibration. (b) Side-view of the breast compression plates, showing the linear translation stage (blue bar on the right) and a linear encoder (in yellow). (c) 3-D rendering of the SLI system, an acrylic bottom plate, and an acrylic compression paddle (top).

### 2.2. Dual-camera SLI breast surface scanning system

The main focus of this report is to characterize and evaluate the SLI-based breast surface acquisition sub-system. This low-profile SLI scanner has a dimension of 30 × 10 × 4.8 cm^3^, and is fixated on a stationary compression plate, on the side facing the patient’s breast [Fig. 2(a)]. It consists of a central projector (P2-B DLP Pico Projector, AAXA Technologies, Irvine, CA, USA) and two USB cameras (C525, Logitech, Lausanne, Switzerland) to reconstruct a 3-D surface of the compressed breast. The SLI scanner is designed to have a relatively short scanner-to-target distance, typically less than 15 cm, and a vertical profile of less than 3 cm to permit scanning breasts with a wide range of sizes. A laptop is used to control the data acquisition, including illumination pattern generation, projection, camera image acquisition, and translation stage control via an interface written in MATLAB (R2017b, Mathworks, Natick, MA, USA).

**Fig. 2.**
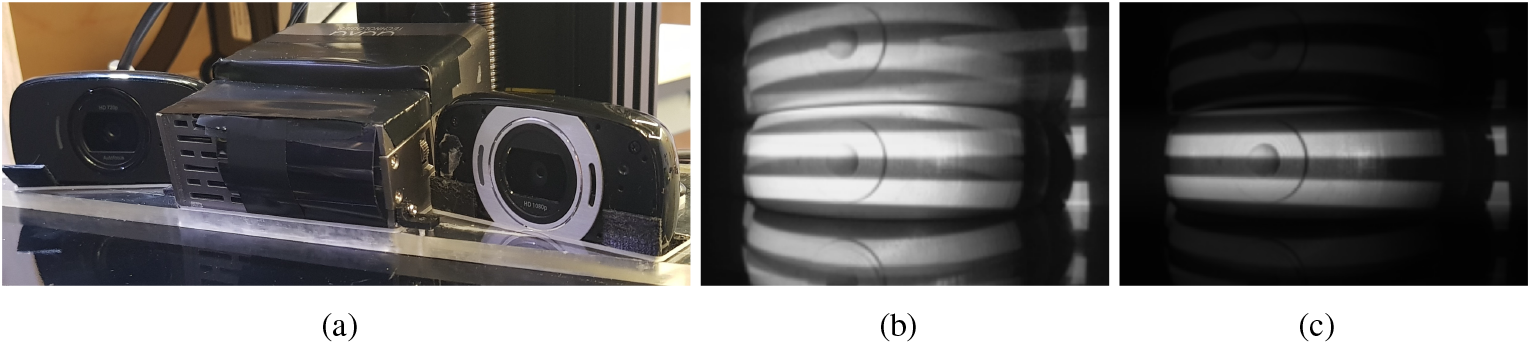
(a) Front-view photo of the SLI system. Cameras and projectors are embedded in an acrylic mount to prevent the need for re-calibration. (b) Original Gray-code pattern showing strong reflection between the compression plates, and (c) cropped pattern informed by separation measurement with brightness level adjusted to avoid saturation.

The principle of SLI is to derive 3-D surface shapes from multiple 2-D images of distorted known patterns [54]. The illumination patterns can be sequential projections, continuous varying patterns, or grid indexing patterns [51]. Calibration is the method of determining the location and orientation of the projector and camera with respect to a global coordinate system [55]. By leveraging this calibration, an SLI system can triangulate multiple light spots on a surface to a location in the projected pattern to derive coordinates of multiple points to generate a point cloud, a collection of 3-D coordinates that represent the surface of the illuminated object [56].

Our SLI system uses complementary Gray-code-based illumination pattern sets due to their robustness to decoding errors [57]. Example Gray-code-based illumination patterns can be found in Fig. 3. Gray-code-based binary patterns [58, 59] are sequentially illuminated onto the breast surface and captured using both USB cameras. These patterns are characterized by their pattern order, *P*. A pattern set of *P* = 3 results in 3 sequences which are a reflected binary of the previous (“01”, “0110”, and “01100110”). Four bar patterns are created for each sequence (a horizontal black and white bar pattern, a vertical black and white bar pattern, and the complementary pattern of each) [60]. The digits correspond to the white (“1”) and black (“0”) bars. In addition, a full-bright (white) and full-dark (black) pattern are added to each pattern set. Thus, a pattern set of *P* = 3 results in 4 × *P* + 2 illumination bar patterns. The two USB cameras have overlapping field-of-views and sequentially capture images of the breast during each illumination pattern at an exposure time of 250 ms. Dual-camera simultaneous acquisition allows the SLI system to capture the curved surface of breasts of varied sizes without moving components.

**Fig. 3.**
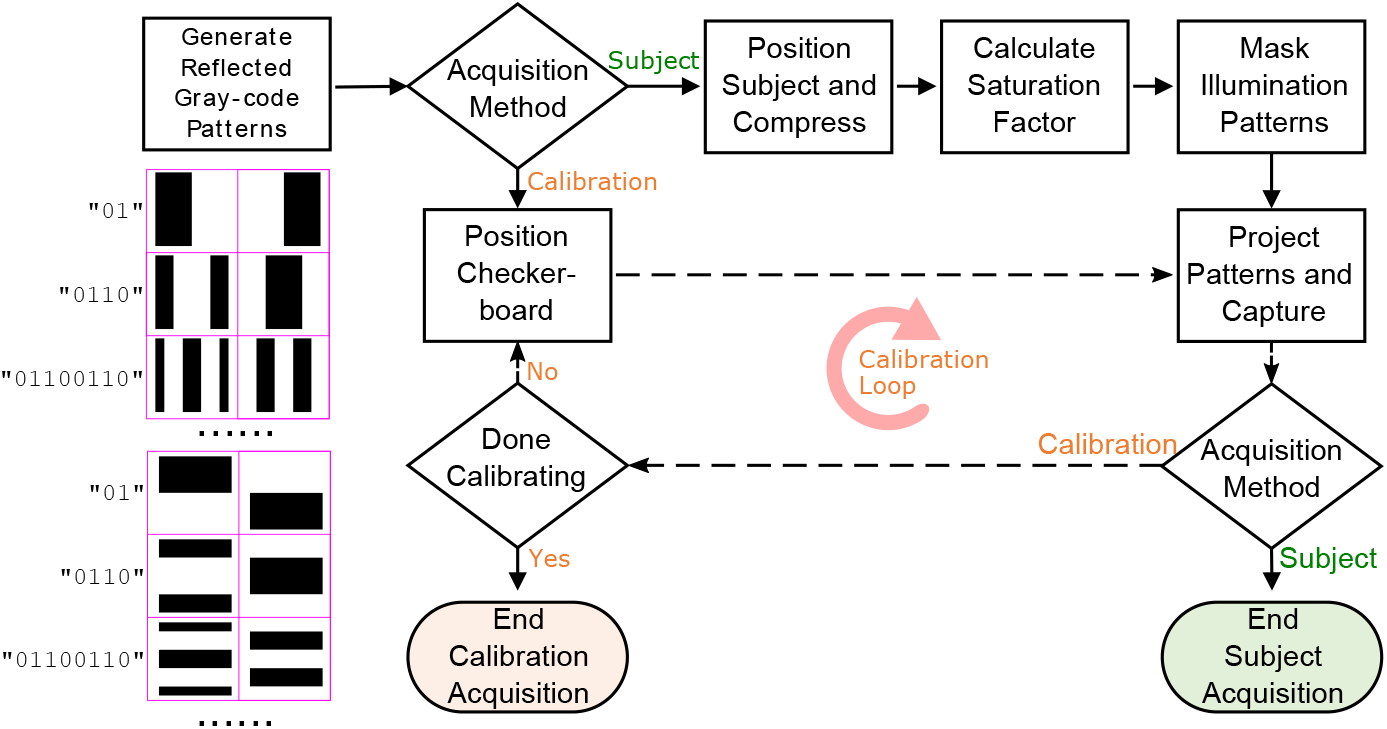
Flow chart of image acquisition for both subject measurements and system calibration. Subject measurements calculate a saturation scaling factor and mask the illumination patterns prior to projecting patterns. System calibration measurements do not mask the illumination patterns and project at full intensity. The calibration loop (dashed lines) is repeated for each location and orientation of the calibration checkerboard. Example Gray-code sequences and the corresponding patterns are shown on the left.

#### 2.2.1. Special data acquisition considerations

While binary-based illumination patterns provide simplicity and robustness towards acquiring surfaces of diverse textures [51], over- and under-exposure of the acquired camera images due to skin colors and projector brightness can impact the quality of the SLI surface. To enhance robustness, the normalized illumination patterns are multiplied by a scaling factor *α* ranging from 0 to 1 to prevent camera saturation and ensure the entire illuminated area can be reconstructed. This ensures we have selected the appropriate illumination brightness that produces captured images with the largest range of pixel values without saturating the cameras. The scaling factor for a camera is calculated prior to data acquisition by first illuminating a full-bright pattern with *α* = 1 onto the breast and capturing a single image using the camera. If the maximum pixel value of the captured image is above a preset threshold, *α* is decreased and the breast is re-illuminated with a full-bright pattern multiplied by the new *α* value. This procedure is repeated until the maximum pixel value of the captured image is less than 95% of the camera’s maximum allowable pixel value. This entire procedure takes an estimated 8 seconds to complete and is repeated for each camera.

Additionally, specular reflections from the acrylic compression plates, shown in Fig. 2(b), can produce vertically mirrored breast surfaces. To minimize such specular reflection, we use dynamic pattern masking based on real-time separation readings provided by a linear encoder. By limiting the vertical span of the illumination patterns, the patterns are projected onto the compressed breast surface without generating strong direct specular reflections from the top and bottom compression plates, as shown in Fig. 2(c). As a result, our SLI system can scan breasts of different compression levels, including breasts without compression.

#### 2.2.2. SLI system calibration and re-projection errors

A standard SLI camera-projector calibration is performed prior to image acquisition and is described in detail in [57]. For each camera-projector pair, a checkerboard pattern is fully illuminated in multiple positions and the corner locations are estimated in the projector’s default coordinate system using a robust pixel classification algorithm [61]. The camera and projector’s intrinsic parameters (optical center and focal lengths) are estimated using a calibration method described in [62] by fixing a world coordinate system to the calibration checkerboard plane.

The projector’s extrinsic parameters (rotation and translation from camera to projector) are calculated using a simple stereo camera calibration [63] that treats the projector as a secondary camera. This results in a rotation matrix and a translation vector relating the camera’s coordinates to the projector’s coordinates. Once the 3-D coordinates of all the corners of the checkerboard are computed using the camera’s (and projector’s) intrinsic and extrinsic parameters, the corners are “reprojected” onto all the images for which they appear. The re-projection error is defined as the average distance between the re-projected corner locations and the actual corner location.

#### 2.2.3. SLI system acquisition

The same acquisition procedures are used for both calibrating the system and acquiring breast shape measurements (Fig. 3). A single acquisition refers to the image capture of all illumination patterns by both cameras. Camera-projector calibration requires an acquisition at each checkerboard position. During breast measurements, the acquisition is preceded by the determination of the saturation scaling factor *α* and masking of the patterns. Patterns during calibration are not masked since the calibration is done with the system fully uncompressed.

### 2.3. Alternative breast surface reconstruction methods for assessing accuracy

To evaluate the accuracy of the SLI system, we compare its output against alternative surface acquisition methods. Each method estimates the surface of a 3-D breast derived from a DBT scan.

#### 2.3.1. Reference breast phantom fabrication

Fig. 4 shows the process of creating surface meshes from DBT scans. Scans were obtained from radiology data from The Cancer Genome Atlas (TCGA) breast Invasive Carcinoma collection [64], available freely through The Cancer Imaging Archive [65]. The scan (ID: TCGA-AO-A03M) was chosen due to its large size and complex surface structure, allowing us to highlight the limitations of low field-of-view acquisition methods and as well as traditional shape estimation methods that simply sweep a single breast contour. Digital Imaging and Communications in Medicine (DICOM) slices were segmented into breast and non-breast regions using ITK-SNAP [66]. Segmented slices were converted to a volumetric image and then into a 3-D mesh using a MATLAB toolbox Iso2Mesh [53] [Fig. 5(a)].

**Fig. 4.**
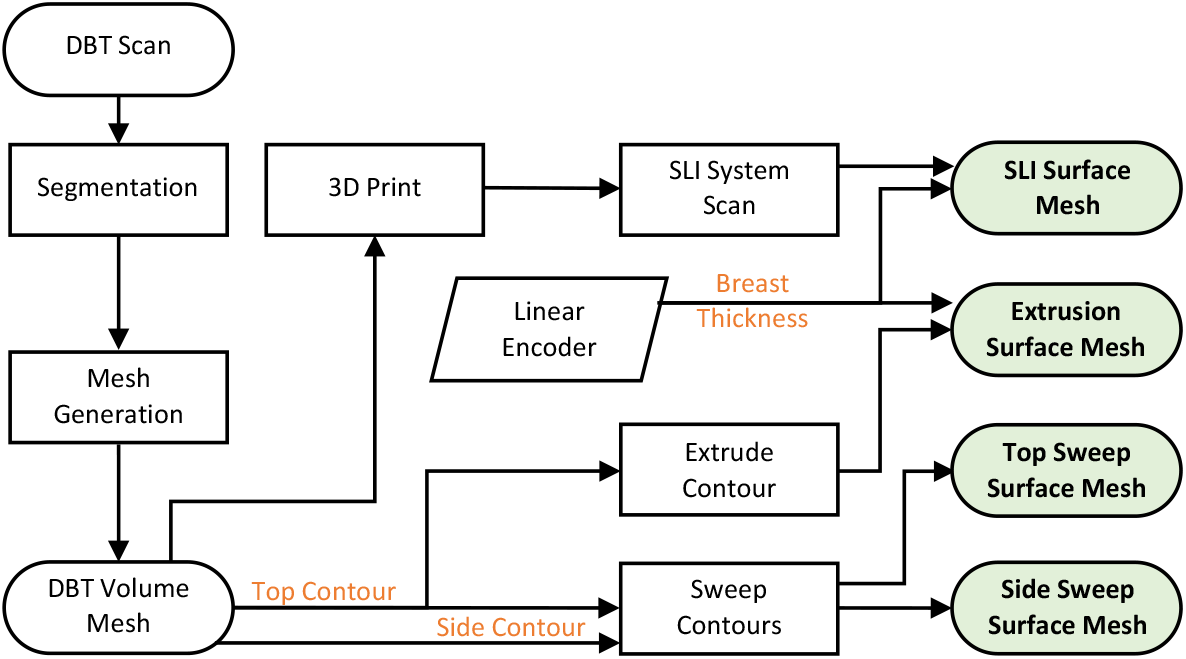
Generation of breast surface meshes using multiple acquisition methods. All mesh models were created based on a DBT volumetric scan.

**Fig. 5.**
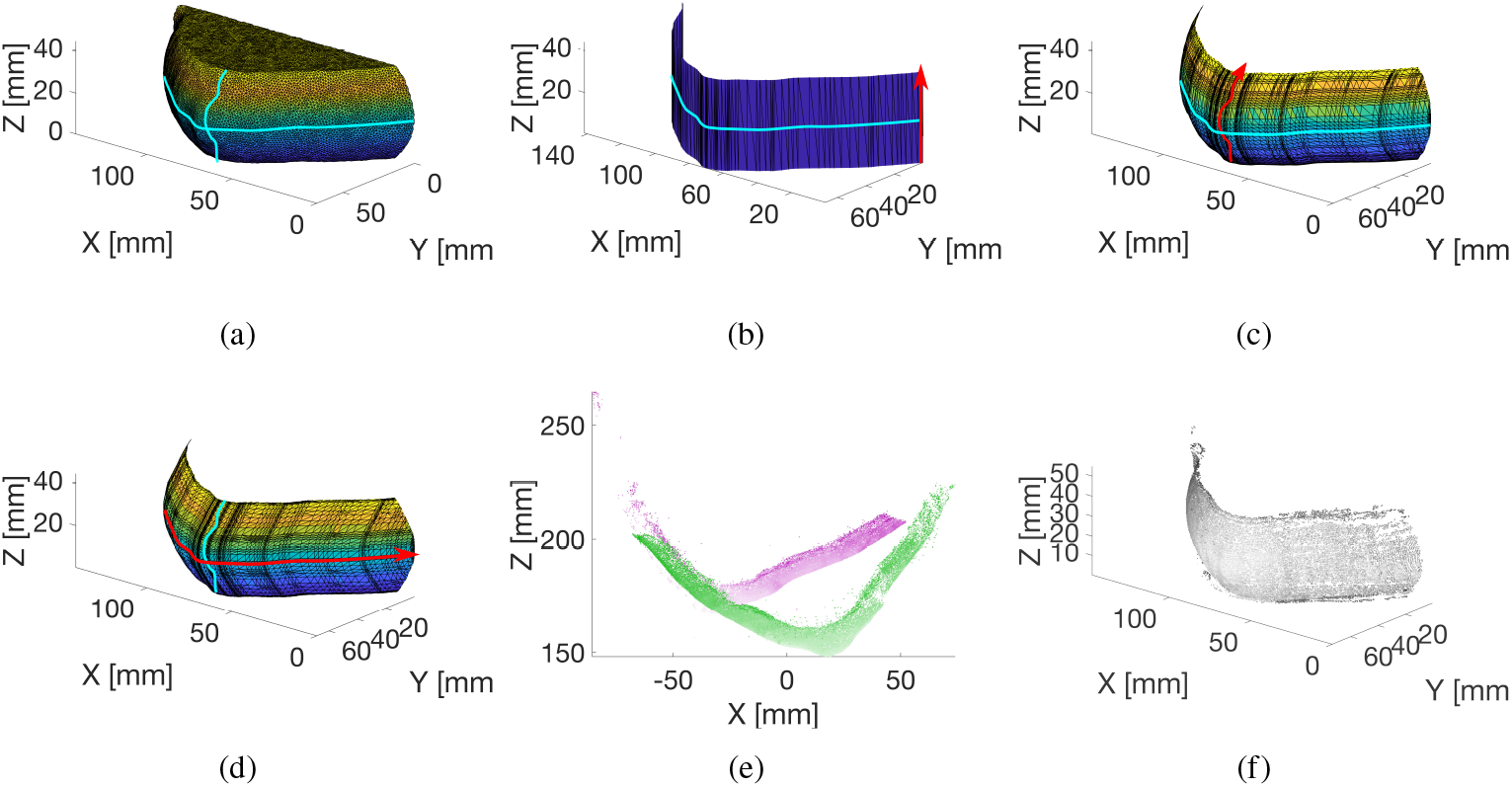
(a) Surface mesh of a digital breast tomosynthesis model obtained from The Cancer Imaging Archive [65]. Blue cyan lines show the *x*/*y* and *y*/*z* breast contours from the top and side views. (b) Estimate of the DBT surface using the extrusion method in which the contour (cyan) is extruded to the thickness of the breast along the *z* axis. (c) The top-sweep method uses the *x*/*y* contour as the profile (cyan) and the *y*/*z* contour as the path to sweep (red). (d) The side-sweep method uses the *y*/*z* contour as the profile (cyan) and the *x*/*y* contour as the path to sweep (red). (e) Point-clouds were generated by scanning a 3-D printed model of the DBT breast using the SLI system. The green (Camera 1) and magenta (Camera 2) point-clouds are in the respective camera coordinates. (f) Merged and denoised point-cloud in the projector’s coordinates.

#### 2.3.2. Single and double contour sweep-based surfaces

Three alternative surface estimation methods are employed in addition to the SLI surface acquisition method. These three methods use spline models of the DBT breast contours from two different planes [cyan lines in Fig. 5(a)]. The extrusion method creates a surface mesh by extruding the *x*/*y* breast contour in the *z* direction to the thickness of the DBT breast measured by the linear encoder [Fig. 5(b)]. The second and third methods utilize a curve-based sweep, in which a profile (shape) follows a path (contour) to create a 3-D model. In the “top-sweep” method, the *x*/*y* contour [cyan line in Fig. 5(c)] is swept along the *y*/*z* contour [red line in Fig. 5(c)]. Similarly, the “side-sweep” method uses the *y*/*z* breast contour [cyan line in Fig. 5(d)] as the profile and the *x*/*y* breast contour as the path [red line in Fig. 5(d)]. In both sweep methods, the profile normal is kept constant.

#### 2.3.3. Structured light imaging surface mesh generation

The SLI system estimates the surface of the compressed breast from the captured images while the breast is illuminated with Gray-code sequence patterns. Each camera-projector pair’s extrinsic parameters are used to generate a point-cloud in each camera’s reference frame using Scan3d-Capture [55] [Fig. 5(e)]. The alignment of each camera-projector pair point-cloud is done by a rigid transformation of each point-cloud to the projector’s coordinates. The point-clouds are then down-sampled using a box grid filter and merged to a single point with normal properties averaged [56]. Denoising is then performed to remove outliers [67]. The point-cloud is trimmed in the *z* direction to the height of the DBT breast measured by the linear encoder [Fig. 5(f)]. The trimmed point-cloud is first converted to a mesh using a crust algorithm [68] prior to being cropped by a bounding-box mesh with height matching the breast thickness to form a closed surface mesh.

#### 2.3.4. Surface estimation error

The surface estimation error, *E*_*s*_, of each surface estimation method is computed by comparing the nodes in each surface mesh to the nodes in the DBT mesh. To obtain the SLI surface mesh of the DBT breast, the DBT breast model was 3-D printed (Ender 5, Creality, China) with a 0.1 mm layer height using white polylactic acid (PLA) filament. The 3-D printed DBT breast was placed in between the compression plates, compressed to the thickness of the printed DBT phantom, and scanned using the dual-camera SLI system.

The residual for each node in the surface mesh is the shortest distance from that node to the DBT mesh. The SLI output mesh is linearly translated (rotation and translation only) into the projector’s frame using the projector’s extrinsic parameters prior to determining residuals. *E*_*s*_ is defined as the average residual of all nodes for a particular surface estimation method.

### 2.4. Evaluation of the impact of surface errors on DOT image reconstructions

Simulations were conducted to evaluate the impact of surface estimation accuracy on DOT reconstruction accuracy for inclusions of various depths. Breast surface meshes were converted to volumetric meshes with optical inclusions and the mean squared error of wide-field DOT reconstructions was calculated for each estimation method.

#### 2.4.1. Assessment of reconstruction accuracy

The effect of different surface estimations on lesion reconstruction was quantified using simulations of continuous wave (CW) pattern-illumination sources. A 5 mm radius spherical inclusion was added at the mid-plane of each volumetric mesh at distances of 5 to 45 mm away from the nipple. The *x* and *z* coordinates of the inclusion were fixed at 68 and 22 mm, respectively. The forward simulation was conducted on a ground truth volumetric mesh consisting of the DBT volumetric mesh and a spherical inclusion. The non-linear image reconstruction of tissue properties was calculated using an iterative Gauss-Newton method in which a series of corrective terms were added to an initial guess [53]. The reconstruction resulted in distributions, 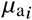, representing the resulting 3-D absorption coefficient (*µ*_*a*_) maps at the *i*^*th*^ node for each simulated tumor location and surface model.

#### 2.4.2. Reconstruction error assessment

We use mean squared error, MSE, to determine the accuracy of the image reconstruction resulting from each breast mesh. To compute the MSE, we first interpolate the reconstructed absorption map, *µ*_*a*_, to the DBT mesh, and then subtract the interpolated *µ*_*a*_ at each node *i*, with the corresponding ground truth absorption value defined on the same node, expressed as

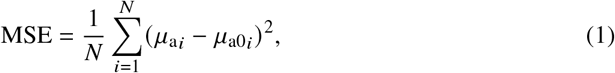

where *N* is the total node number; 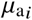 and 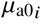 define the recovered and ground truth *µ*_*a*_ values, respectively, at the *i*^*th*^ node in the DBT mesh. A lower MSE value indicates higher accuracy.

## 3. Results

In this section, we first report the projector and camera re-projection errors of our SLI calibration using the calibration checkerboard. We then quantify the error of surface estimation methods in estimating the surface shape of the DBT breast. Finally, we show the effect of different surface estimation methods on optical property reconstruction using simulations of continuous wave pattern-illumination sources.

### 3.1. Camera-projector calibration and surface acquisition

Our dual-camera SLI system was calibrated in a dark room using a checkerboard with 5 × 7 internal corners with 1 × 1 cm^2^ black and white squares. The calibration checkerboard was printed and adhered to a black Delrin surface to ensure it remained planar. To account for varying breast shapes and curvatures, the checkerboard was placed at 7 locations. At each location, camera images were captured for 5 board orientations: 1) normal to the *y*-axis [see Fig. 1(a)], 2) rotated left and 3) rotated right by 30 degrees relative to the *x*-axis, and 4) tilted forward and 5) tilted backward by 30 degrees in the *y*/*z* plane [Fig. 1(b)]. This results in a total of 7 × 5 = 35 checkerboard positions within the camera and projector field-of-views (Fig. 1). Each rotation and tilt was measured manually using a printed protractor. The projector’s resolution is 1280 × 720 pixels and the resolution of the cameras is 1600 × 896 pixels. Using a Gray-code of bit-length *P* = 9, we acquire *P* × 4 + 2 = 38 images (see Section 2.2) at each board orientation/position placement. An exposure time of 0.25 seconds per image per camera results in a total one-time calibration time of 38 × 7 × 5 × 2 × 0.25 = 665 seconds. The first camera-projector pair (Camera 1 with projector) resulted in a camera and projector re-projection error of 0.4089 and 0.2282 pixels, respectively. The second camera-projector pair resulted in a camera re-projection error of 0.4368 pixels and a projector re-projection error of 0.2889 pixels.

A re-calibration is only necessary when the relative position of the cameras and projector is changed. Once calibrated, the SLI system can acquire a surface scan in about 35 seconds, including 16 seconds for adaptively adjusting the intensity scaling factor *α* for both cameras (see Section 2.2.1 for details) and 19 seconds for image acquisition (38 × 2 × 0.25 = 19 s).

### 3.2. Surface estimation errors

A basic breast model was created to highlight our system’s ability to capture surface shape features. This model is extruded by a breast contour derived from Fig. 5 in [27]. We added a 25.4 mm diameter cylindrical extrusion to represent an areola and a 12.7 mm diameter semi-sphere representing a nipple. This model was then 3-D printed with its surface acquired using our SLI system. Fig. 6 shows the resulting point-cloud overlaid on the basic breast model. Not only does the point-cloud follow the large contour of the 3-D printed basic breast model, but it also captures the geometry of the areola and nipple [Fig. 6(b)]. The surface estimation error, *E*_*s*_, between the point-cloud and the original surface was 0.2479 mm. The reconstructed areola yielded a diameter of 25.78 mm.

**Fig. 6.**
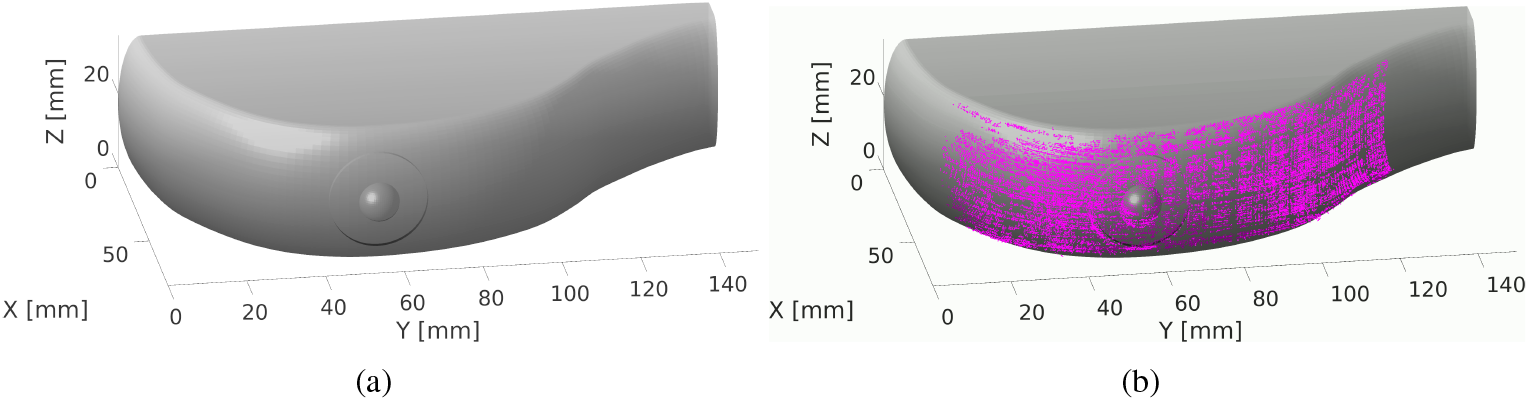
Validation of the SLI scanner using a breast model containing basic shapes. In (a), we show the surface model used for 3-D printing of the phantom; in (b), we overlay the SLI-reconstructed point-cloud (magenta) over the original model.

Additionally, the SLI system calculated a saturation scaling factor of *α* = 0.8 for both cameras when the 3-D printed digital breast tomosynthesis (DBT) breast was scanned. The number of nearest neighbor points was set to four and the outlier threshold was set to one standard deviation from the mean of the average distance to those four neighboring points. The resulting point-cloud from the SLI system scan has 35,256 points.

Table 1 shows the mean and standard deviation of the residual of all the nodes in the estimated breast surface mesh. The *z*-extrusion method (EXT) results in the largest error (*E*_*s*_) of all compared methods. While the top-sweep, side-sweep, and SLI methods all had similar standard deviations, the SLI method resulted in the smallest *E*_*s*_.

**Table 1.** Mean and standard deviation of the residuals of each point in a surface estimation mesh compared to the original DBT breast mesh.

|  | Extrusion | Top-Sweep | Side-Sweep | SLI |
| --- | --- | --- | --- | --- |
| Surface estimation error, $E_s$ [mm] | 6.8353 | 0.3772 | 0.4726 | 0.2543 |
| Standard deviation [mm] | 2.8671 | 0.3029 | 0.3370 | 0.2723 |

### 3.3. Mean square error of optical property reconstruction

DOT reconstructions were performed using our in-house data analysis toolbox, Redbird-m [69]. An *L*-curve analysis is used to determine the Tikhonov regularization parameter as 3. × 16 10^−10^, which is fixed over 10 Gauss-Newton iterations. The absorption coefficient of the spherical inclusion was set to be twice (*µ*_*a*_ = 0.016/mm) that of the background tissue (*µ*_*a*_ = 0.008/mm). The reduced scattering coefficient *µ*^′^_*s*_ was set to 1 mm^−1^ for both breast and inclusion tissues. A set of 32 (16 vertical, 16 horizontal) moving-bar source patterns [70] covering an area of 40 × 40 mm^2^ was centered at the spherical inclusion. Iso2Mesh was used to interpolate nodal values from the reconstructed mesh to the ground truth mesh based on linear interpolation in order for all reconstructed meshes to have the same number of nodes.

The MSE errors from these reconstructed images are summarized in Fig. 7, showing the effect of different surface estimation methods on the accuracy of optical property recovery. Overall, surface mesh accuracy appears to have a notable impact on relatively shallow tumors, with a depth of less than 25 mm. Although MSE values obtained using the SLI method closely follow those using the ground truth DBT mesh, they remain higher than DBT MSE values for all inclusion depths, indicating a less accurate reconstruction. The top- and side-sweep-based meshes followed similar trends, however, reporting higher errors compared to SLI especially when the tumor is relatively shallow. The maximum MSE value for the SLI mesh at a distance of 5 mm from the surface (4. × 89 10^−7^ mm^2^) was 23% higher than the maximum MSE value for the DBT mesh (4.35 × 10^−7^ mm^2^). In contrast, the single-axis-extrusion method (EXT) MSE was nearly twice higher (8.62 × 10^−7^ mm^2^) than that from the DBT mesh. Although the DBT and SLI mesh MSEs plateau to their minimum around 15 mm from the surface, top-, side-, and extrusion-based mesh MSEs continue to decrease until a depth of 25 mm. Beyond the depth of 25 mm, the errors between different methods become minimal.

**Fig. 7.**
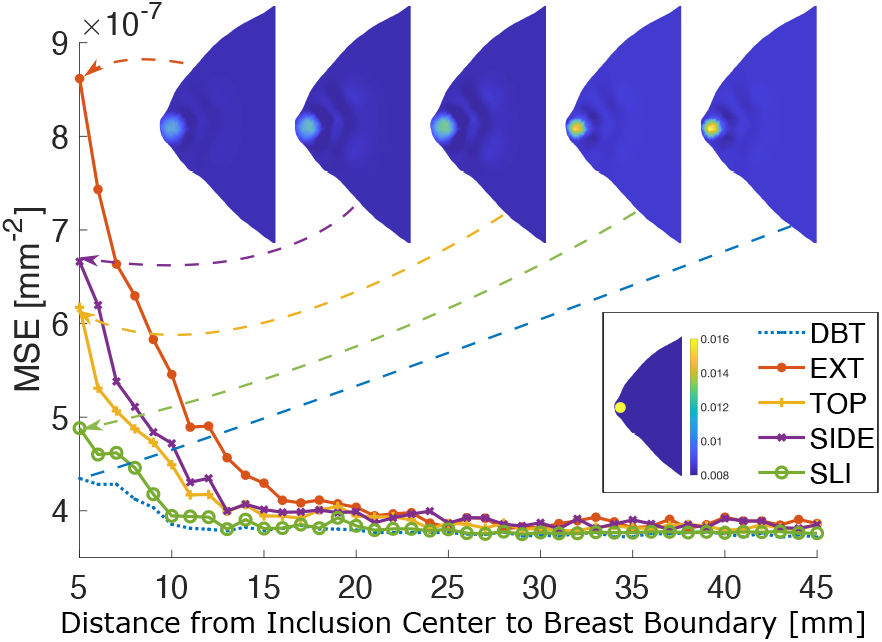
A comparison between the mean squared error (MSE) of the reconstructed absorption map using 4 estimated surfaces (EXT -*z*-axis extrusion, TOP – sweeping *x*/*y* contour along *y*/*z* contour, SIDE – sweeping *y*/*z* contour along *x*/*y* contour, and SLI – surface acquired from our SLI system) as well as the ground truth surface (DBT). A 1 cm diameter spherical inclusion is moved away from the breast surface at various depths between 5 and 45 mm in 1 mm increments. Image slices (in *x*/*y* plane) of the reconstructed absorption coefficient (*µ*_*a*_ in mm^−1^) (top-row) and the ground truth *µ*_*a*_ (top-right) are shown as insets.

## 4. Discussion

The camera and projector re-projection errors in Section 3.1 represent an average error of less than 0.5 pixels in estimating the corner locations of a calibration checkerboard placed between 50 and 250 mm away [Fig. 1(b)] from the projector for all 35 checkerboard positions. Although the same illumination patterns and calibration checkerboard positions were used to calibrate each camera-projector pair, we find a slightly better calibration accuracy (a re-projection error difference of 0.06 pixels) when the projector is paired with Camera 1 since Camera 1 is closer to the projector’s lens (Fig. 1). This small discrepancy in the re-projection errors of the two pairs is due in part to the asymmetry of the dual-camera setup. The asymmetry arises from the projector offset relative to its housing, making one camera closer to the projector than the other [Fig. 1(a)]. Although independent camera-projector pair calibrations compensate for the asymmetry of the dual-camera setup, other factors that influence camera-projector calibration accuracy are hardware arrangement and the number of input images used for calibration [71]. Our system addresses hardware arrangement by maximizing the common viewing area between the projector and cameras by placing them as close as possible. We also affix the projector and cameras to an acrylic base to minimize movement, which minimizes the variability of the calibration results. Additionally, we use 38 images for calibration, well above the suggested 15 calibration images after which the re-projection error stabilizes [71].

From Table 1, the single-axis extrusion method resulted in the highest surface error because it does not account for the curvature of the breast in the *y*/*z* plane [Fig. 5(b)]. Table 1 indicates that, on average, points in the extrusion-method-derived surface estimation mesh are approximately 6.84 mm away from the DBT mesh. The top- and side-sweep methods decrease the surface estimation error by incorporating a second breast contour from the *y*/*z* plane [Figs. 5(c) and 5(d)]. Both methods improve the accuracy of surface estimations by approximating the 3-D curvature of the breast. We want to point out that both top-sweep and side-sweep methods require an additional camera to obtain two orthogonal views of the breast [72], which does not necessarily lead to simplified hardware compared to the SLI setup considering the mounting space constraints and lighting conditions [52]. While also requiring two cameras, our mammography-tailored SLI system can produce sub-millimeter resolution of the surface compared to the reference DBT breast model based on Table 1.

Our results also demonstrated that the improvement in surface estimation accuracy can lead to improved DOT reconstruction accuracy. Fig. 7 shows using breast surfaces derived from SLI can accurately recover the absorption profile compared to those recovered using the ground-truth (DBT) mesh at most tested tumor depths. For superficial/shallow (< 10 mm) tumors, the top- and side-sweep surface estimation methods followed similar trends to each other, reporting MSEs about 50% higher compared to those from using ground-truth (DBT) surface models, and about 30% higher than those from using SLI surfaces. As expected, the effect of the surface accuracy decreases as the inclusion is moving further away (> 25 mm) from the skin.

Despite the ability to produce sub-millimeter resolution of breast surfaces in poorly lit and confined mammography-like settings, both our SLI system and our analysis have limitations. Firstly, the span of the output point-cloud from our SLI system is limited to the area of the breast that is well-illuminated by the projector. As a result, tissue boundaries near the chest wall or those in direct contact with the compression plate may not be well covered due to the limited angles of the projector/camera line-of-sight. Still, for DOT of a compressed breast, capturing a significant portion of the front-facing breast tissue as our system does, provides quantitative differences in reconstructions, as shown above. Future improvement of this system should consider using more compact, wide-angle projectors, higher resolution cameras, and patterns with higher order binary codes to both expand the field-of-view and increase the point-cloud resolution. Secondly, a 3-D printed breast model was used to experimentally compare different shape acquisition methods. Different choices of extruder sizes, filament colors, and printing techniques could impact the surface texture of the printed phantom and slightly alter the surface estimation errors. Thirdly, our adaptive intensity scaling method is used to prevent over- and under-exposure to achieve more robust surface reconstructions. Finally, the quantification of reconstruction errors was based on simulations using a single set of pre-determined breast models, tumor size and shape, tumor contrast, and wide-field pattern size. An experimental validation using heterogeneous phantoms may produce more realistic comparisons.

## 5. Conclusion

In summary, we have developed and validated a low-profile, low-cost, and robust SLI-based breast surface acquisition system that can be used in confined low-light mammography-like settings to obtain 3-D breast surfaces. Once calibrated, our SLI system can achieve sub-millimeter accuracy with a data acquisition time of less than 40 seconds. We quantified the impact of breast surface estimation methods on DOT optical property reconstruction accuracy of inclusions embedded at various depths and found that obtaining accurate breast surfaces is important for DOT reconstructions of shallow lesions with a depth less than 25 mm. While contour-extrusion based approaches are relatively simple and produce acceptable reconstructions for deeply embedded tumors, they can result in 30% to 100% higher errors when reconstructing shallow tumors. We want to particularly mention that a compact breast shape acquisition system that can fit between mammography compression plates can not only help improve parallel-plate breast DOT image reconstructions but can also be incorporated into standard x-ray based DBT scanners to help improve 3-D DBT image reconstructions. Currently, clinical DBT image reconstructions are performed without considering the actual breast shape [73] and often result in an inaccurate cylindrical quasi-3D breast shape [69]. Explicitly capturing and considering breast 3-D shapes are expected to lead to improved image quality in DBT and other model-based breast imaging modalities.

